# Fallopian tube precursor lesions of serous ovarian carcinoma require L1CAM for dissemination and metastasis

**DOI:** 10.1101/270785

**Authors:** Kai Doberstein, Rebecca Spivak, Yi Feng, Sarah Stuckelberger, Gordon B. Mills, Kyle M. Devins, Lauren E. Schwartz, Marcin P. Iwanicki, Mina Fogel, Peter Altevogt, Ronny Drapkin

## Abstract

Most high-grade serous ovarian carcinomas (HGSC) arise from Serous Tubal Intraepithelial Carcinoma (STIC) lesions in the distal end of the fallopian tube (FT). FT secretory cells become malignant by accumulating genomic aberrations over 6-7 years before seeding the ovarian surface, with rapid tumor dissemination to other abdominal structures thereafter. It remains unclear how nascent malignant cells leave the FT to colonize the ovary. This report provides evidence that the L1 cell adhesion molecule (L1CAM) contributes to the ability of transformed FT secretory cells (FTSEC) to detach from the tube, survive under anchorage-independent conditions, and seed the ovarian surface. L1CAM was highly expressed on the apical surface of STIC lesions and contributed to ovarian colonization by upregulating integrin and fibronectin in malignant cells and activating the AKT and ERK pathways. These changes increased cell survival under ultra-low attachment conditions that mimic transit from the FT to the ovary. To study metastasis to the ovary, we developed a tumor-ovary co-culture model. We showed that L1CAM expression was important for FT cells to invade the ovary as a cohesive group. Our results indicate that in the early stages of HGSC development, transformed FTSECs disseminate from the FT to the ovary in L1CAM-dependent manner.

The authors have declared that no conflict of interest exists

**List of abbreviations:**

FT: fallopian tube
FTSEC: fallopian tube secretory epithelial cell
HGSC: high-grade serous carcinoma
IHC: immunohistochemistry
OSE: ovarian surface epithelium
STICS: Serous Tubal Intraepithelial Carcinomas
TCGA: The Cancer Genome Atlas
AKT: Protein kinase B
ERK: extracellular-signal-regulated kinase

## Introduction

Ovarian cancer, the most lethal gynecologic malignancy in developed countries, is a heterogeneous disease with multiple histologic subtypes (Kroeger and Drapkin 2017) that accounts for nearly a quarter of a million incident cases and over 150,000 deaths worldwide each year (Ferlay et al. 2015; Siegel et al. 2017). The most common subtype of ovarian cancer is high-grade serous carcinoma (HGSC) and most patients with HGSC eventually develop recurrent disease that is resistant to cytotoxic chemotherapy. The poor survival rates are in part due to the lack of early detection tools and late clinical diagnosis (Karst and Drapkin 2010; Vaughan et al. 2011). Despite these dire statistics, significant progress has been made in recent years in our understanding of the pathogenesis of HGSC. Based on histologic, molecular, and genetic evidence, it is now generally accepted that a majority of HGSC arise in the secretory cells of the distal fallopian tube (FT) (Kurman and Shih Ie 2011; McDaniel et al. 2015; Eckert et al. 2016; Labidi-Galy et al. 2017; Meserve et al. 2017). More recently, next generation sequencing of fallopian tube precursors showed that mutations in *TP53* are among the earliest genetic events, occurring in benign appearing secretory cells that are non-proliferative. Stretches of these *TP53* mutated secretory cells are called ‘p53 signatures’ and are considered benign in isolation. The acquisition of malignant cytological features and proliferation with expansion of these secretory cells leads to the formation of Serous Tubal Intraepithelial Carcinomas (STICs). Using whole-exome sequencing and mathematical modeling, we showed that the average time between development of a STIC lesion and ovarian cancer is approximately 6.5 years (Labidi-Galy et al. 2017). Unfortunately, once the malignant FT cells encamp on the surface of the ovary, seeding of the peritoneal cavity occurs rapidly thereafter. These data suggest that the ability of malignant FTSECs to disseminate and metastasize to the ovary is a critical event during the development of HGSC. It is currently not known what are the molecular events that trigger FT lesion to metastasize to the ovary.

The L1 cell adhesion molecule (L1CAM) is a type-1 transmembrane molecule that is over-expressed in various types of human cancers, including HGSC (Altevogt et al. 2016) and may have an important role in neoplastic processes (Bondong et al. 2012). Many studies have shown that the expression of L1CAM is associated with malignant characteristics such as chemoresistance, epithelial to mesenchymal transition (EMT), proliferation, migration, invasion and survival (Kiefel et al. 2012). In HGSC, it was shown that increased expression of L1CAM is associated with worse overall survival and may confer chemoresistance in this setting (Fogel et al. 2003; Bondong et al. 2012; Doberstein et al. 2014). L1CAM is expressed in the majority of HGSCs and its expression, as well as the presence of soluble L1CAM in ascites fluid, contributes to the invasive and metastatic properties of HGSC (Fogel et al. 2003; Bondong et al. 2012). It is not known at what stage of ovarian cancer development L1CAM is expressed and whether or not it has a functional role in this process.

In this report, we provide evidence that L1CAM plays a critical role in transformation and dissemination of HGSC FT precursors to the ovary. We found that L1CAM was highly expressed on the apical surface of STIC lesions and that its expression contributed to the dissemination of cells. L1CAM promoted FTSEC sphere formation and survival of ovarian cancer cells and immortalized FT cells. L1CAM mediated this effect through the activation of the Integrin, AKT and ERK pathways which increased cell survival when cultured under substrate detachment conditions. Furthermore, utilization of a 3D *ex-vivo* model of human FTSEC spheres and mouse ovaries revealed that L1CAM was important for FT cells to invade the ovary as a cohesive group. Our results provide important mechanistic insights about the early steps of ovarian cancer metastasis from the fallopian tube to the ovary.

## Results

### L1CAM expression and prognosis in high grade serous ovarian carcinoma

To determine the expression of L1CAM in HGSC, we analyzed L1CAM RNAseq expression from The Cancer Genome Atlas (TCGA) data set. We first compared L1CAM expression in HGSC to other cancers from the Pan-Cancer cohort, a dataset containing over 10,000 different tumor samples (Figure 1A) (Cancer Genome Atlas Research et al. 2013). Our analysis revealed that HGSC exhibits some of the highest levels of L1CAM, second only to gliomas and skin cancer (Cancer Genome Atlas Research 2011). We then analyzed the association between L1CAM expression and overall survival. Using the Kaplan Meier plotter (kmplot.com), we found that high L1CAM expression is strongly associated with reduced overall survival (Supplementary Figure 1A), a finding consistent with previous studies (Bondong et al. 2012; Doberstein et al. 2014).

**Figure 1:**
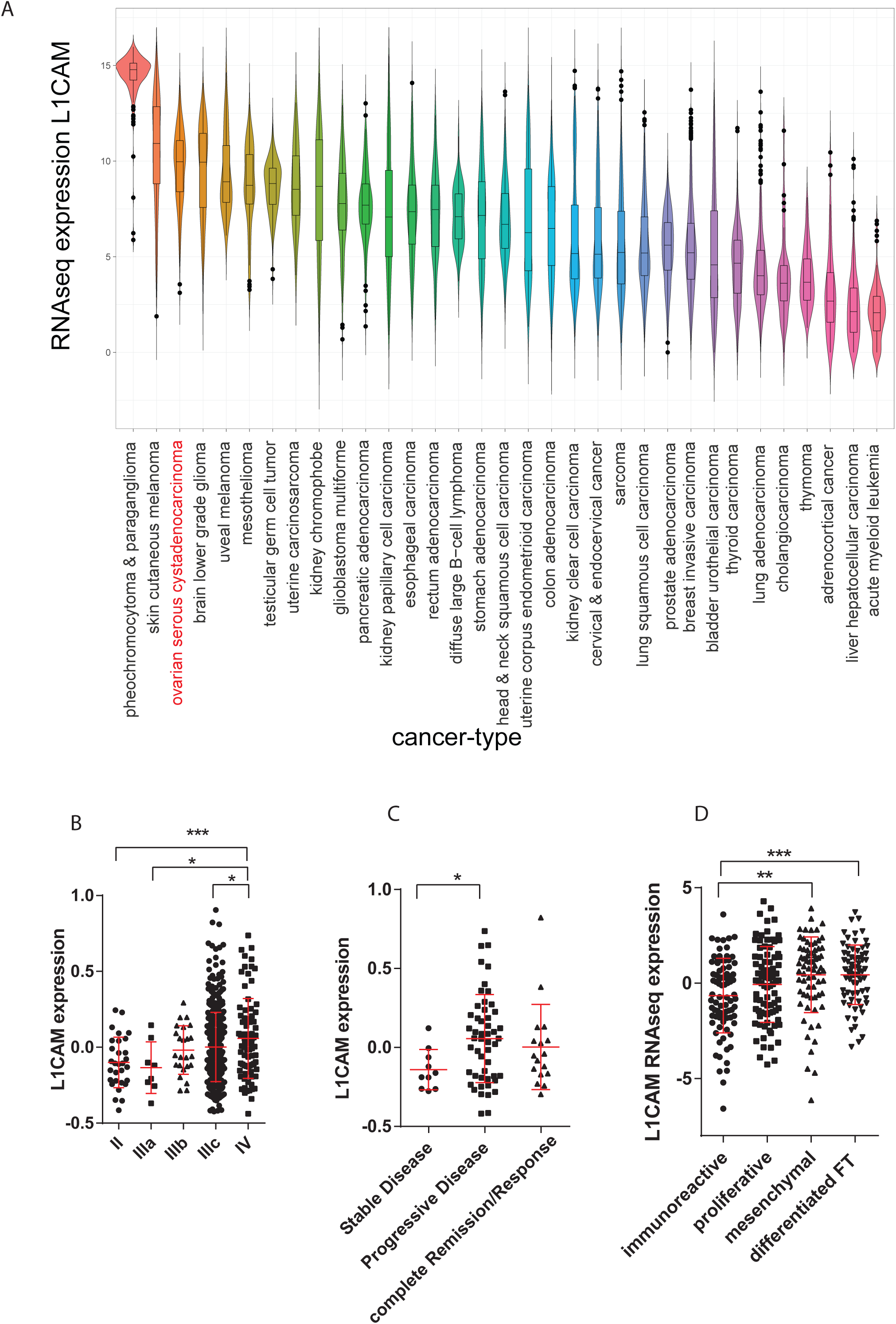
L1CAM expression in the TCGA ovarian cancer cohort: (A) Violin plot of L1CAM expression in different cancer types from the PAN-Cancer cohort. (B) L1CAM mRNA expression in tumors of the TCGA ovarian cancer cohort in different histological stages. (C and D) The same samples were analyzed for the relationship of L1CAM expression to disease progression and TCGA cluster. P-values were calculated with an unpaired two-sided t-test. *=p<0.05, **=p<0.01, ***=p<0.001.

The correlation of L1CAM expression with survival suggests that L1CAM may contribute to disease progression. Using TCGA RNAseq data, we found that L1CAM expression exhibits a gradual increase from Stage II to Stage IV disease (Figure 1B). Interestingly, the biggest increase occurred between Stage IIIA and Stage IIIC. These stages differ by the degree of metastatic spread beyond the pelvis to the peritoneal cavity or omentum. Additionally, we found that L1CAM is expressed significantly higher in patients with progressive disease compared to those with stable disease (Figure 1C). When analyzing L1CAM expression in the four different molecular subgroups described in the original TCGA study (immunoreactive, proliferative, mesenchymal and differentiated), we found the highest L1CAM expression in the mesenchymal and differentiated fallopian tube-like subgroups; notably the mesenchymal subgroup displays the worst prognosis (Cancer Genome Atlas Research 2011) (Figure 1D). Overall, those data support previous findings showing that high L1CAM expression correlates with poor patient outcomes (Doberstein et al. 2014) and highlight its putative role in ovarian carcinoma progression.

### Expression of L1CAM in precursor lesions of the fallopian tube

To determine whether L1CAM plays a role in early stages of serous ovarian carcinoma, we analyzed its expression by immunohistochemistry (IHC) in fallopian tube precursor lesions. Immunohistochemistry for p53, p16, and Stathmin was used to credential seventeen cases of FT specimens with STIC lesions as previously described (Karst et al. 2011b; Novak et al. 2015). Using a L1CAM specific antibody (Fogel et al. 2014; Doberstein et al. 2015), we found that L1CAM expression was readily detectable in 16 of the 17 STIC lesions (Figure 2A, 2B). Interestingly, we did not observe homogenous protein expression throughout the STIC lesions, but rather a gradient with peak expression near the apical regions of the STIC lesions (Figure 2A, 2B and 2C). Some of the cells that stained strongest for L1CAM also appeared to be detached from the main STIC lesion, suggesting that L1CAM is enriched in disseminating STIC lesions (Bijron et al. 2013). In support of this hypothesis, in one section we found P53-and L1CAM-positive cell clusters that were detached from the original lesion and detected in an adjacent region of normal fallopian epithelium (Figure 2D). We further confirmed that this pattern was independent of p53 status as we observed a similar expression in STIC lesions that were negative for p53 expression (Figure 2A). In 10 benign FT epithelial sections we observed only occasional L1CAM-positive cells (Figure 2E). Together, these data suggest the hypothesis that L1CAM is involved in the regulation of FTSEC dissemination from the fallopian tube.

**Figure 2:**
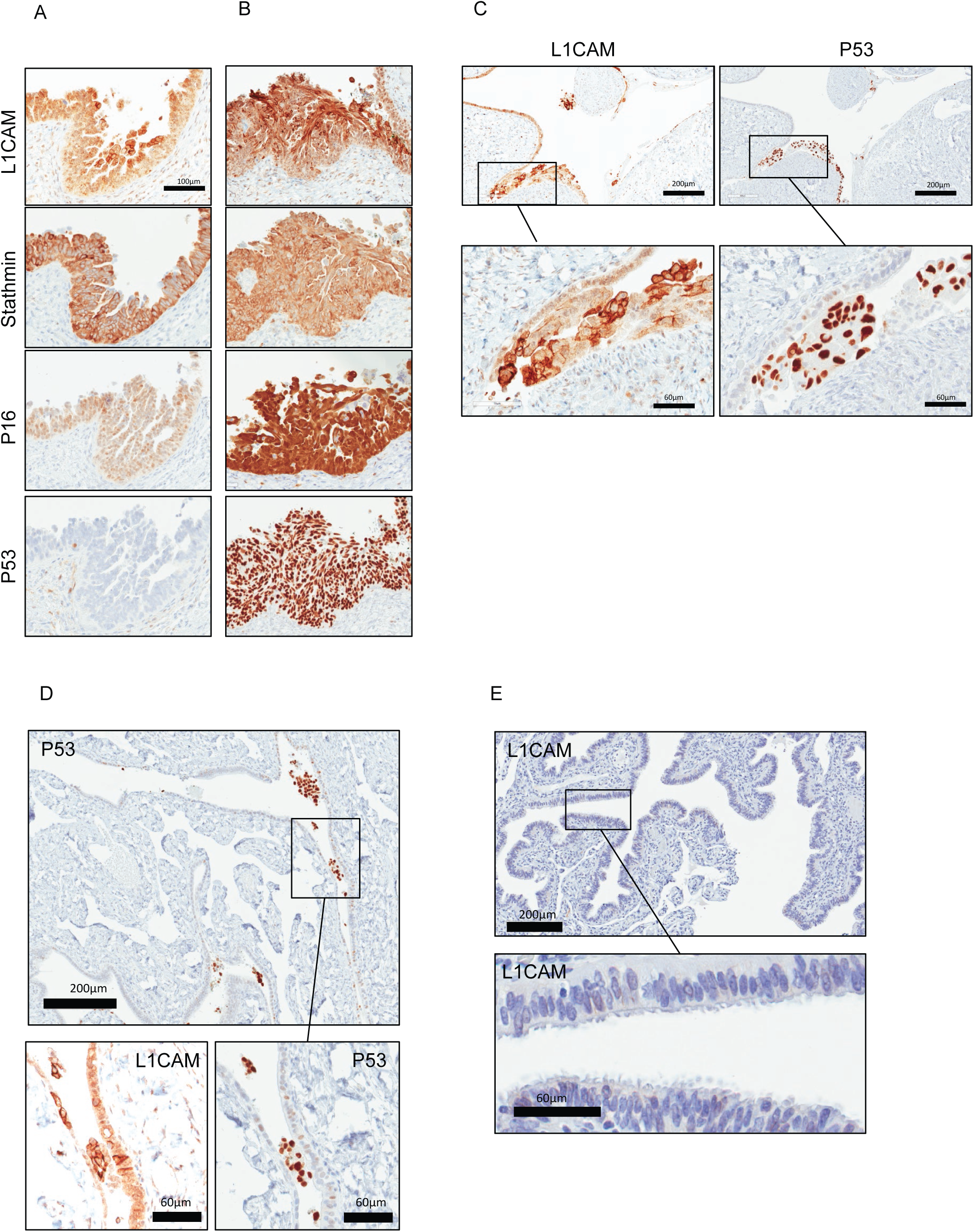
L1CAM expression in STIC lesions: (A) P53-negative STIC lesion and (B) P53-positive STIC lesion. Both were stained for L1CAM, Stathmin, P16, and P53. C) STIC lesion stained for L1CAM and P53. (D) Normal fallopian tube tissue with P53-positive metastatic cells floating in the tubal lumen. (D, E, F) Scalebar represents 100 μm.

### Characterization of L1CAM in ovarian cancer and fallopian tube cells

To begin deciphering the role of L1CAM in HGSC pathology, we analyzed L1CAM expression in a panel of ovarian cancer cell lines by Western blot and found that all of the cell lines expressed detectable L1CAM protein (Figure 3A). Similarly, in primary cells derived from ascites fluid of HGSC patients, we found high L1CAM expression in seven of nine samples (Figure 3B). In primary FTSECs that were derived from healthy donors and immortalized using different techniques (Karst et al. 2011a; Karst and Drapkin 2012), we found that some cell lines displayed little or no expression (FT33, FT190, FT237 and FT246), some showed moderate expression (FT189, FT194, FT240 and FT282) and one showed high expression (FT282CE) of L1CAM (Figure 3C). The heterogeneous pattern in L1CAM expression among the samples likely reflects the genetic diversity amongst different individuals and the effect of different immortalization techniques during FTSEC culture establishment. Immunofluorescence for surface bound-membranous L1CAM correlated with our Western blot data (Figures 3D-3F). The frequent expression of L1CAM in these cells suggested to us that it may have a role in ovarian cancer. Since migration plays an important role during the progression and dissemination of ovarian cancer, we wanted to evaluate the role of L1CAM in ovarian cancer cell lines and engineered FTSEC. We used RNA interference to evaluate the impact of L1CAM knockdown on a panel of L1CAM-expressing cell lines. We observed a significant reduction in migration in all lines tested (Figure 3G, Supplementary Figure 1C). Conversely, when we overexpressed L1CAM in fallopian tube cell lines that showed low or moderate expression of L1CAM, we found a significant increase in cell migration (Figure 3H). These data support the hypothesis that L1CAM is functionally involved in the regulation of ovarian cancer dissemination from the fallopian tube.

**Figure 3:**
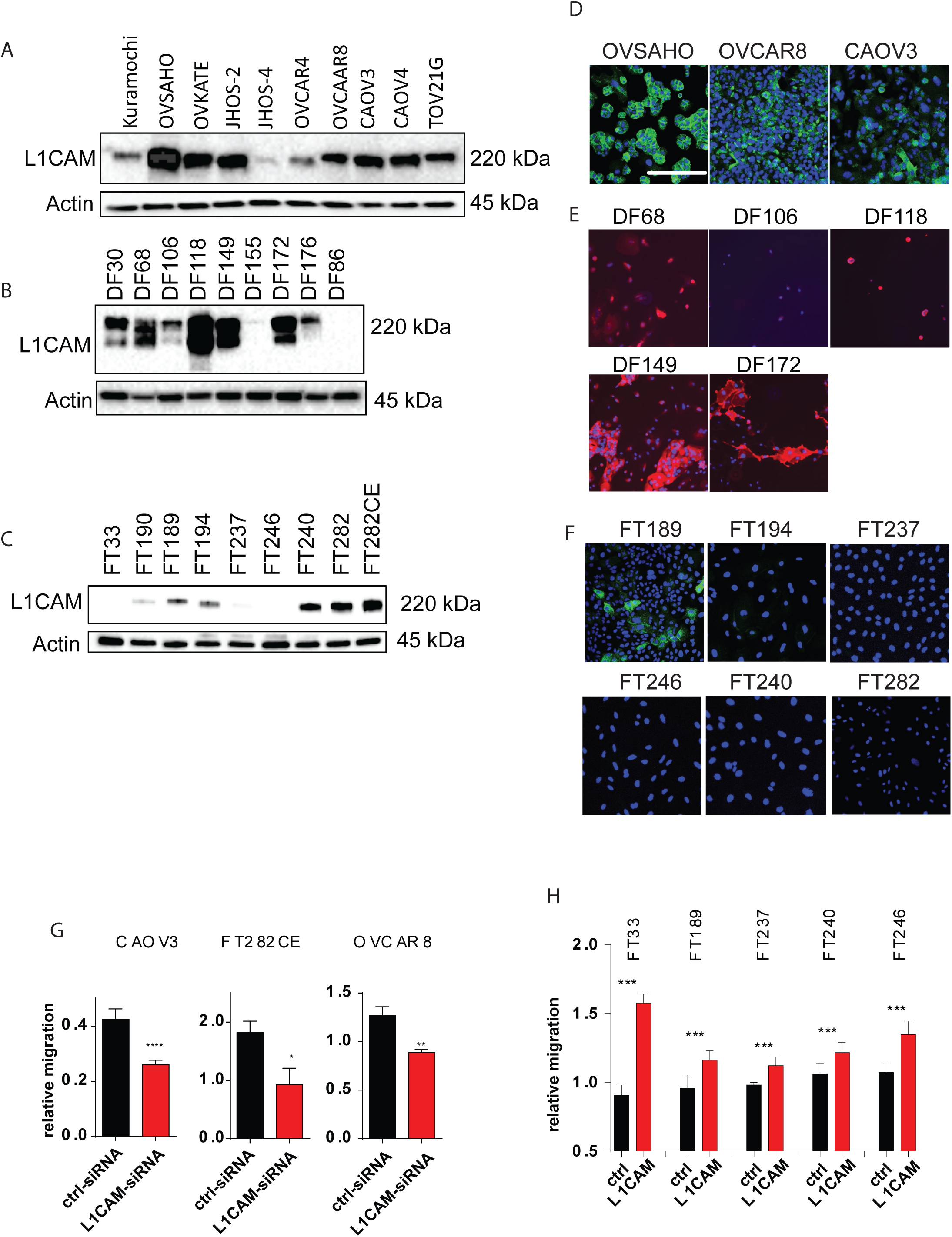
L1CAM expression in Ovarian cancer and primary cell lines: (A, B and C) Western blot analysis of L1CAM expression in ovarian cancer cell lines (A), primary ascites-derived HGSC cells (B) and immortalized fallopian tube cell lines (C). Actin was used as a loading control. (D, E, F) Immunofluorescence images of selected cell lines for L1CAM with DAPI to stain the nucleus. (G) Relative migration in CAOV3, FT282-CE and OVCAR8 after knock-down of L1CAM. (H) Relative migration measured after overexpression of L1CAM or the empty control vector. (D, E, F) Scalebar represents 100 μm. Three independent experiments were performed and P-values were calculated with an unpaired two-sided t-test. *=p<0.05, **=p<0.01, ***=p<0.001, ****=p<0.0001.

### Knock-out of L1CAM by CRISPR/Cas9 inhibits sphere formation in OVCAR8 cells

Another characteristic of ovarian cancer dissemination from the fallopian tube to the ovaries is the ability of tumor cells to form free floating multicellular structures. To examine whether L1CAM might help to trigger the cellular dissemination from STIC lesions, we used the OVCAR8 cell line that spontaneously forms multicellular structures in culture. We noticed that when OVCAR8 was allowed to grow for an extended period of time on standard adherent plates, the cells gained the ability to transition from a two-dimensional (2D) monolayer to three-dimensional (3D) cell structures that ultimately detached (Figure 4A, Supplementary Figure 1D and E). In addition, OVCAR8 cells were also able to form spheres when grown under ultra-low attachment (ULA) conditions. Since this process of coalescence and subsequent detachment of cell clusters observed *in vitro* showed similarity to the L1CAM-expressing cell clusters observed in STIC lesions (Figure 2), we speculated that L1CAM might be involved in the process of sphere formation. Therefore, we used CRISPR/Cas9 to knockout L1CAM in OVCAR8 cells (Figure 4B). These cells were termed OVCAR8-ΔL1 cells. We found that the loss of L1CAM strongly reduced their ability to form colonies and impeded their ability to invade (Figure 4C and 4D, respectively). The L1CAM-negative cells showed a reduced ability to form spheres under 3D conditions, which was most pronounced after 1 and 2 days, but was less dramatic after six days in culture (Figure 4F), indicating an important role of L1CAM for the initiation of sphere formation. The loss of L1CAM also reduced compaction of the spheres, which was measured as change in size (Figure 4E). Similarly, when grown on an adherent surface for one week, the OVCAR8-ΔL1 cells continued growing in a monolayer, whereas the L1CAM-positive wildtype cells built 3D structures (Figure 4A). Importantly, when grown under serum free conditions, the ability to build spheres was completely abrogated in the OVCAR8-ΔL1 cells (Figure 4G) leading to an increase in cell death, as measured by uptake of ethidium bromide, in those cultures (Figure 4G and 4H). We could partially rescue the ability of the cells to build spheres by overexpressing L1CAM in the OVCAR8-ΔL1 cells (Figure 4I and 4J).

**Figure 4:**
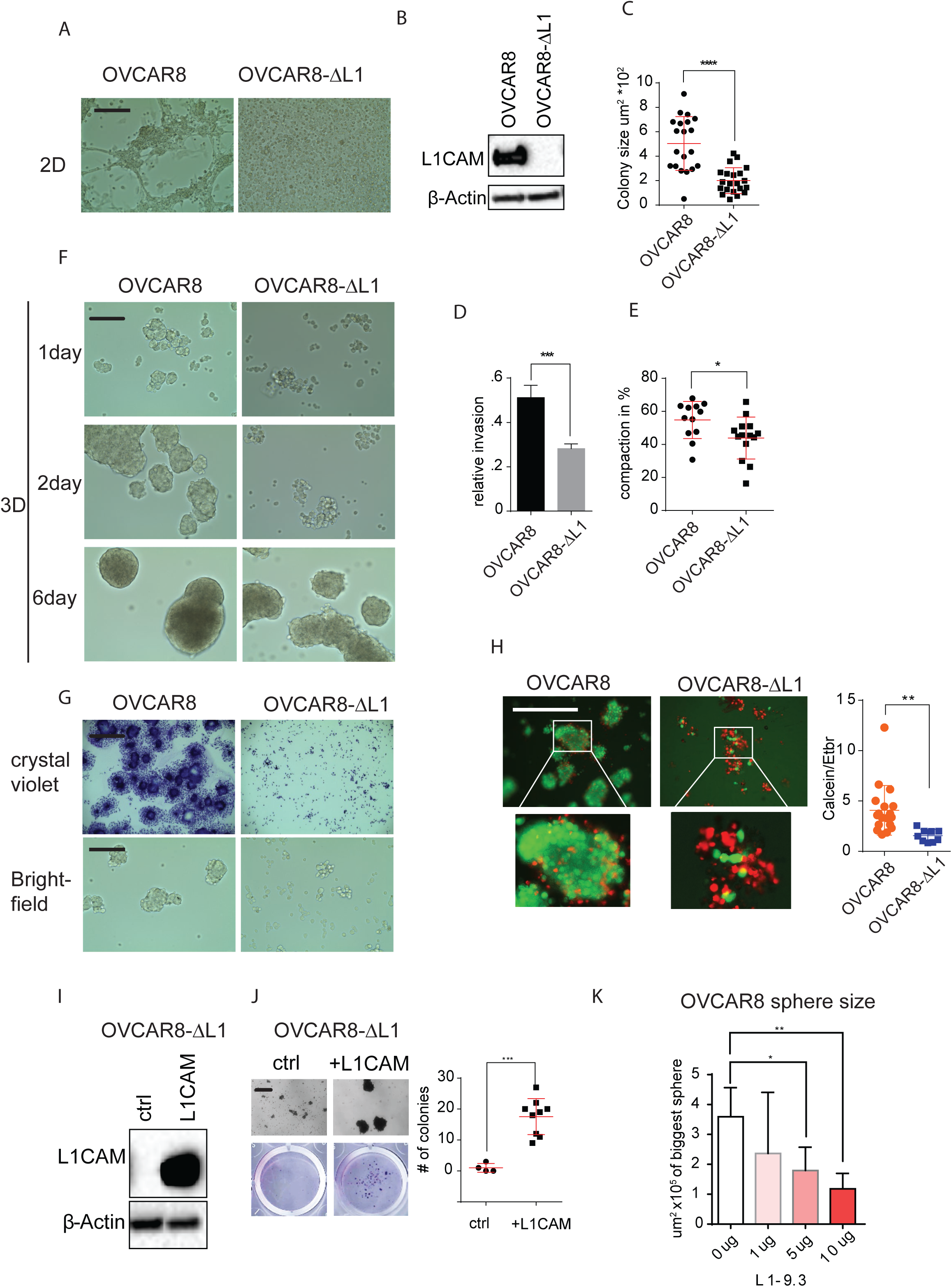
L1CAM knockdown inhibits sphere formation in OVCAR8 cells: (A) phase contrast images representing OVCAR8 and OVCAR8-ΔL1 cells grown for 6 days under 2D conditions. (B) Western blot analysis of L1CAM expression in wild type (wt) OVCAR8 and OVCAR8-ΔL1 cells after the transient transfection of CRISPR/Cas9 with a guide RNA targeting L1CAM. (C) Colony formation assay was quantified for colony size in different OVCAR8 cell populations. (D) Relative invasion of OVCAR8 and OVCAR8-ΔL1 cells. (E) Cells were measured for compaction (area change). (F) phase contrast images representing OVCAR8 and OVCAR8-ΔL1 cells grown for 1, 2 and 6 days grown under 3D conditions and for 6 days. (G) OVCAR8 and OVCAR8-ΔL1 cells analyzed by bright field microscopy (lower row) and for 3D colonies with crystal violet (upper row) after culturing under 3D conditions in serum free media. (H, left panel) Fluorescent images representing OVCAR8 and OVCAR8-ΔL1 stained for calcein (green) and ethidium bromide (red). (G, right panel) Dot plot showing the ratio between living and dead cells, assessed by measuring the ratio between calcein (green) and ethidium bromide (red) intensity in OVCAR8 and OVCAR8-ΔL1 cells after culturing for 7 days under 3D and serum free conditions. (I and J) Rescue experiment: Re-expression of L1CAM analyzed by western blot (I) in OVCAR8-ΔL1 and its effect on sphere formation (J) Dot plot representing number of colonies after cells were transfered from 3D to 2D cell culture. (A, F, G, H, J) Scalebar represents 200 μm. (K) Effects of L1CAM antibody treatment (anti-L1-9.3) on sphere size after 48 hours. P-values were calculated with an unpaired two-sided t-test. *=p<0.05, **=p<0.01, ***=p<0.001, ****=p<0.0001

To address the question whether the extracellular domain of L1CAM contributed to sphere formation, we examined whether a L1CAM-specific antibody (clone L1-9.3), that can block L1CAM homophilic binding (Wolterink et al. 2010), could disrupt this process. Increasing concentrations of the antibody led to a partial reduction in sphere size (Figure 4K). Interestingly, the effect of the antibody was most pronounced during the early phase of sphere formation, concordant with the effects of the L1CAM knock out. If spheres were allowed to form, the antibody had only a modest effect on sphere stability.

### Overexpression of L1CAM in primary fallopian tube cells increases sphere forming ability

To examine whether L1CAM is able to drive dissemination of HGSC precursors from the FT, we asked whether the overexpression of L1CAM could trigger a similar effect in immortalized fallopian tube cell lines as seen in OVCAR8 cells. Specifically, we overexpressed L1CAM in the FT cell lines FT237, FT240, FT246 and FT282 and compared them to the parental cells expressing an empty vector control (Figure 5A). When seeded on a 2D surface we detected an increase in 3D structures and colonies that were comparable to the structures observed in OVCAR8 cells. In contrast, the L1CAM-negative FT cell lines maintained a monolayer (Supplementary Figure 2A). Similarly, when seeded on ULA plates, we found a marked increase in sphere formation in L1CAM-overexpressing FT237, FT240, FT246 and FT282 cells, as analyzed by sphere size (Figure 5B and 5C). The increase in sphere formation led to a reduction in cell death and an increase in the number of living cells when FT237 and FT240 cells were cultured under ULA and serum-free conditions (Figure 5D and Supplementary Figure 2B, 2C, respectively). Importantly, when cultured in the presence of the L1CAM blocking antibody L1-9.3, sphere size was significantly smaller compared to the IgG control (Figure 5E), but sphere formation was not prevented, consistent with our findings with OVCAR8. These results indicate that L1CAM contributes to sphere formation of fallopian tube epithelial cells.

### L1CAM-positive cells form heterotypic spheres under ULA culture conditions

The observation in STIC lesions and OVCAR8 cells that L1CAM leads to the formation of multicellular spheres, defined by robust cell compaction and adhesion, led us to hypothesize that cells expressing high levels of L1CAM might preferentially interact with other cells that express high levels of L1CAM. To test this hypothesis, we overexpressed L1CAM in two different fallopian tube cell lines and co-cultured these cells at a ratio of 1:10 with non-transduced control cells under ULA conditions. To visualize the co-cultures, we labeled the L1CAM-overexpressing FT cells red and the control cells green with fluorescence dyes. Interestingly, similar to the expression in STIC lesions, the L1CAM overexpressing cells clustered together within the FT sphere (Figure 5F). When we co-cultured OVCAR8 with normal fallopian tube cell lines at a ratio of 1:10, the OVCAR8 cells clustered and attached as cell groups to the fallopian tube clusters, but did not intersperse into the fallopian tube cells (Figure 5G). However, when co-culturing the OVCAR8-ΔL1 cells with the fallopian tube cells, we observed a stronger intermixing of both cell types (Figure 5G). These data indicate that cells expressing high levels of L1CAM bind preferably to other L1CAM expressing cells rather than to cells lacking L1CAM.

**Figure 5:**
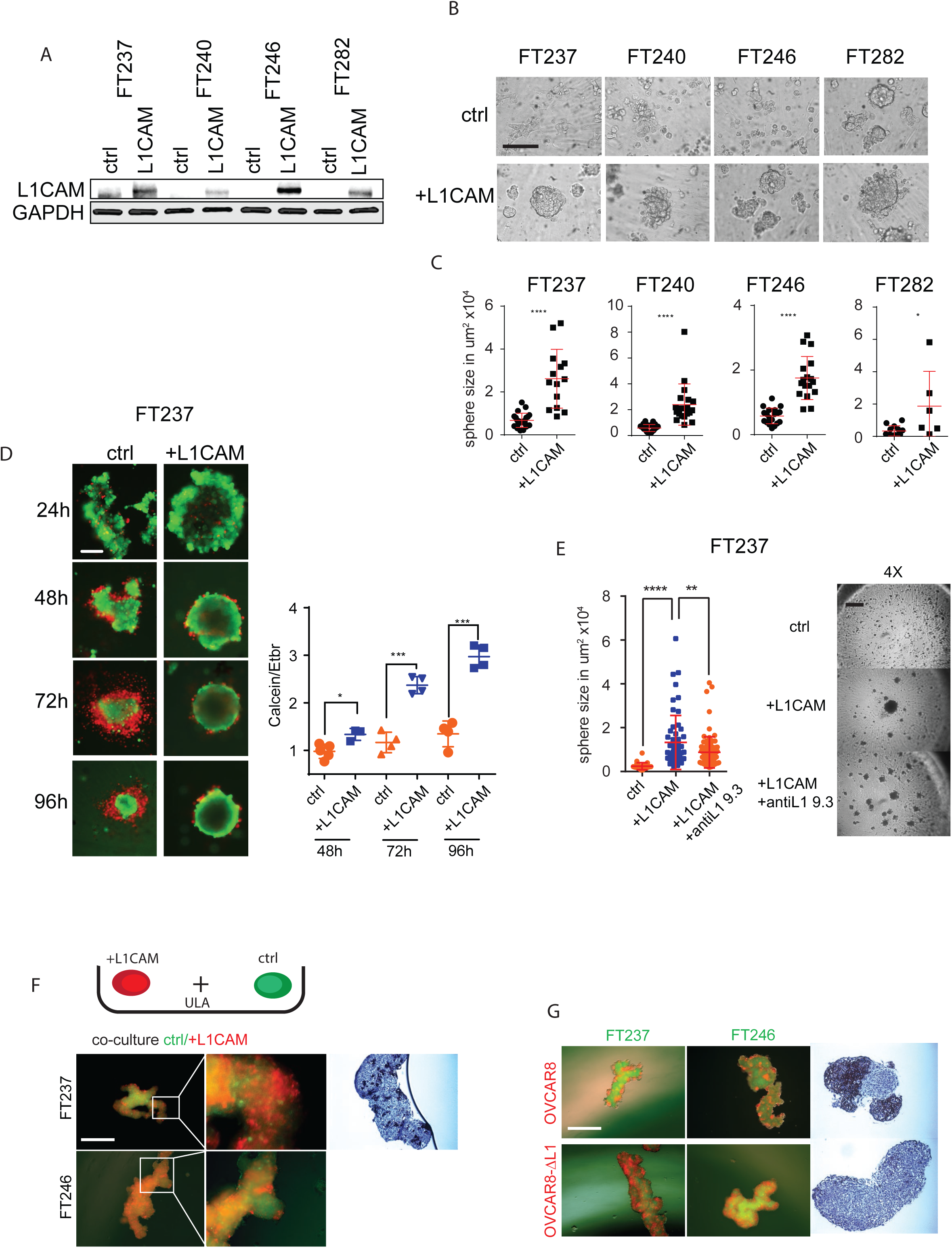
L1CAM triggers sphere formation in primary fallopian tube cell lines: (A) Western blot analysis of primary fallopian tube cell lines transfected with L1CAM or a control vector and analyzed for L1CAM and GAPDH (housekeeping gene) expression. (B) Bright field images of primary fallopian tube cell lines from A. Scalebar represents 200 μm. (C) Quantification of area in um^2^ of spheres shown in B. (D) (right panel) Fluorescent images representing FT237 cells transfected with a L1CAM or (left panel) a control vector. Dot plot depicting the ratio between living and dead cells, assessed by measuring the ratio between calcein (green) and ethidium bromide (red) intensity in FT237 cells transfected with L1CAM or a control vector after culturing under 3D and serum free conditions. Scalebar represents 100 μm. (E, left panel) Dot plot showing sphere size of FT237 cells that was analyzed after transfection with control vector or L1CAM with the addition of either IgG antibody or the anti L1-9.3 antibody. (E, right panel) Phase contrast of cells transfected with control vector (left) or L1CAM with the addition of either IgG antibody or the anti L1-9.3 antibody. Scalebar represents 250 μm. (F) schematic cartoon showing the co-culture experiment under ultra low adhesion conditions (ULA). Fluorescent images of co-culture of FT237 (upper row) or FT246 (lower row) cells that were transfected either with L1CAM (red) or a control vector (green) and co-cultured at a ratio of 1:10, respectively. (G) Fluorescent images of co-culture of OVCAR8 or OVCAR8-ΔL1 (red) with either FT237 or FT246 (green) in a ratio of 1:10, respectively. (F, G) Scalebar represents 500 μm.

### L1CAM expression promotes extracellular matrix deposition and integrin expression to form spheres in immortalized fallopian tube cells under 3D culture conditions

To analyze which pathways were altered in OVCAR8 versus OVCAR8-ΔL1 cells cultured under 2D and 3D conditions, we performed a reverse phase protein array (RPPA) analysis (Supplementary Figure 3A). One of the most dramatic changes we noted was the near complete loss of fibronectin expression in the L1CAM-negative cells grown in 2D and 3D conditions (Figure 6A, Supplemental Figure 3A). However, growth of L1CAM-wildtype OVCAR8 cells under 3D conditions led to an increase in fibronectin expression that was confirmed by Western blot analysis (Figure 6A, Supplemental Figure 3A). Previous studies have documented the important roles of fibronectin and Integrin-α5β1 in the establishment of extracellular matrix support for ovarian cancer cells (Iwanicki et al. 2016). Consistent with these functions, we observed a strong downregulation of Integrin-α5β1 in the OVCAR8-ΔL1 cells compared to OVCAR8 cells, whereas integrin-β3 was upregulated in OVCAR8-ΔL1, indicating a change in the integrin signaling pattern.

**Figure 6:**
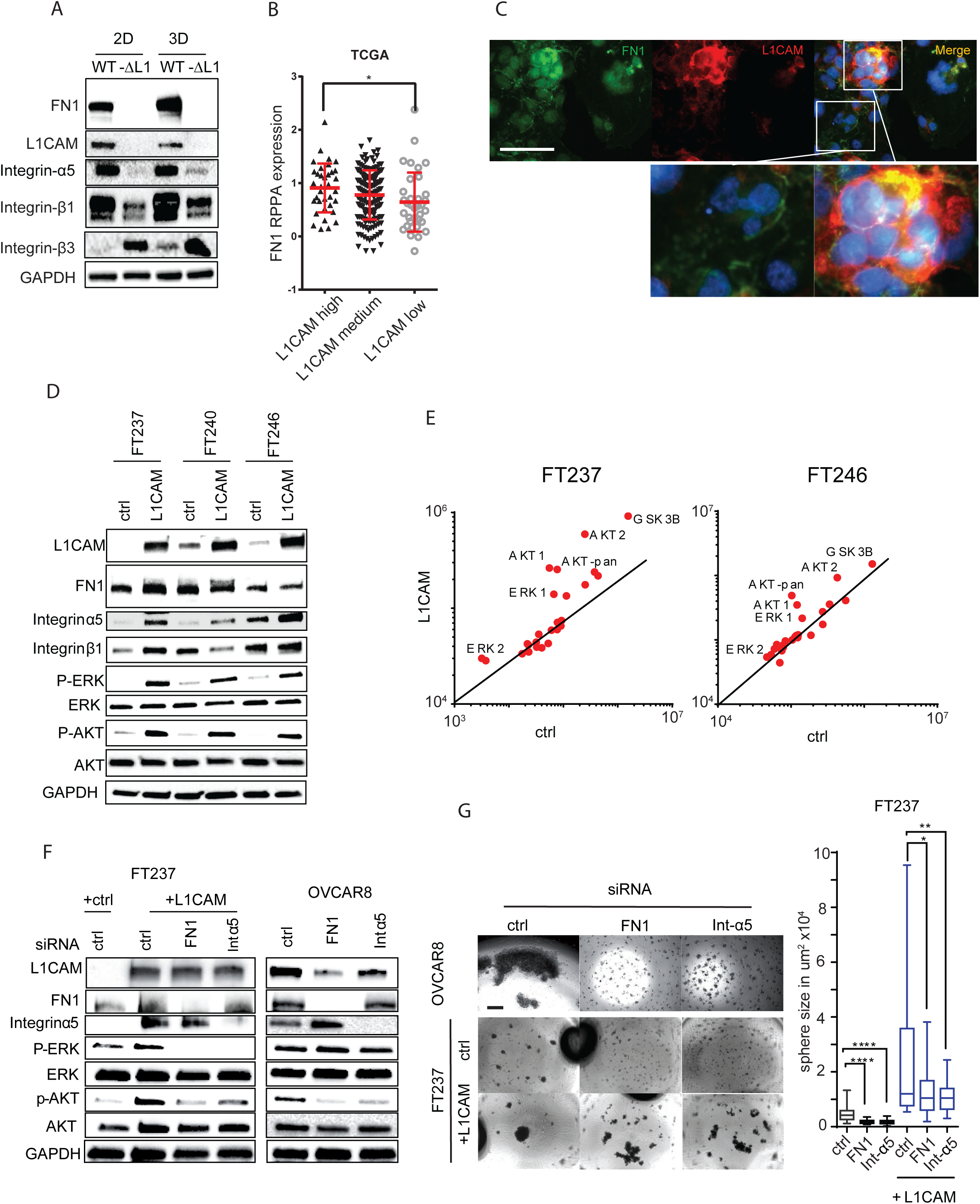
L1CAM regulates the expression of fibronectin during 3D sphere formation: (A) Western blot analysis of fibronectin, Integrin α5, Integrin β1, Integrin β3, L1CAM and GAPDH in OVCAR8 and OVCAR8-ΔL1 under 2D and 3D culturing conditions. Representative experiment (n=5) is shown (B) Dot plot representing fibronectin RPPA protein expression in ovarian cancer tumors of the TCGA cohort with a high, medium and low L1CAM mRNA expression. (C) Fluorescent images representing L1CAM (red) and fibronectin (green) in OVCAR8 cells cultured under 3D conditions before re-seeding on coverslips for fixation. Scalebar represents 200 μm. (D) Western blot analysis of FT237, FT240 and FT246 transfected with either L1CAM or a control vector. Representative Western blot (n=3) is shown. (E) Scatter plot representing protein kinase array in FT237 and FT246 expressing control plasmid and plasmid-containing L1CAM. (F) Western blot analysis of FT237 and OVCAR8 analyzed 72h after siRNA treatment, targeting fibronectin, Integrin α5. Representative Western blot (n=3) is shown (G) Left: Bright field images representing OVCAR8, FT237-ctrl and FT237+L1CAM cells grown under ULA conditions in serum free media, 72h after siRNAs-mediated attenuation of fibronectin or Integrin α5 expression. Scalebar represents 400 μm. Right: Box plot representing quantification of sphere size of FT237-ctrl and FT237+L1CAM after siRNA knockdown of fibronectin or Integrin α5.

We then analyzed the protein expression of fibronectin (Figure 6B: RPPA-data, Y-axis) in the ovarian cancer TCGA dataset. We split the cases into three groups based on their L1CAM mRNA (X-axis) expression (high, medium and low; determined relative to median expression) and found that tumors expressing high levels of L1CAM also expressed significantly more fibronectin protein (Y-axis) (Figure 6B).

To analyze the co-expression of L1CAM and fibronectin under 3D and 2D culture conditions, we re-seeded the spheres overnight on a cover slide and analyzed the attached spheres by immunofluorescence for fibronectin and L1CAM. Interestingly, we found that L1CAM and fibronectin showed the highest co-expression at the interface of the sphere, while cells that attached to the plastic surface grew in a monolayer and showed a much lower expression (Figure 6C). We next analyzed whether overexpression of L1CAM in primary FT cell lines also regulates the expression of fibronectin and Integrin-α5β1 (Figure 6D). While all FT cell lines already exhibited fibronectin expression, L1CAM overexpression caused a further increase in fibronectin expression, most notably in FT237. Additionally, we observed an increased expression of both Integrin-α5 and integrin-β1 subunit in all the cell lines. These data indicate that L1CAM promotes the expression of fibronectin and Integrin-α5β1 in primary FT cells.

### Overexpression of L1CAM activates the ERK and AKT pathway in fallopian tube cells

Our data indicate that L1CAM is required for the expression of integrin α5β1 and fibronectin; cell adhesion molecules that have been implicated in ovarian cancer cell survival and dissemination (Burleson et al. 2004; Mitra et al. 2011; Scalici et al. 2014). In addition, L1CAM expression has been reported to activate mitogen-activated protein kinase (MAPK) pathways in several cell line models (Silletti et al. 2004; Gast et al. 2005). Thus, we hypothesized that L1CAM is involved in the regulation of pro-survival signaling pathways in transformed FTSEC and cancer cell lines. To test our hypothesis, we analyzed a MAPK and AKT array after overexpression of L1CAM in two immortalized FT cell lines. We found a strong activation of ERK and AKT1/2 in the L1CAM-expressing cells compared to controls (Figure 6D and 6E, Supplementary Figure 3C). Correspondingly, we observed a reduced activity in both pathways in the OVCAR8-ΔL1 cells (Supplementary Figure 3B), which is also in agreement with previous reports (Gast et al. 2005; Doberstein et al. 2011; Nakaoka et al. 2017). Furthermore, siRNA-mediated attenuation of fibronectin or integrin-α5 expression abrogated the activation of ERK and AKT in the fallopian tube cell lines, while it decreased AKT but not ERK activation in OVCAR8 cells (Figure 6F). Importantly, the knockdown of fibronectin or Integrin-α5 also reduced the ability of OVCAR8 cells to form spheres (Figure 6F-6G), suggesting that cell-matrix-cell adhesion mediated by integrin α5 supports tumor cell survival. These results are consistent with the role of integrins and ECM in supporting ovarian cancer dissemination.

### L1CAM is sufficient to support ovarian cancer colonization of the ovary

Our results support the hypothesis that L1CAM promotes FTSEC dissemination from the fallopian tube to the ovary, through the stabilization of integrin/matrix complexes that activate pro-survival pathways including AKT and ERK. To test whether ovarian cancer cells and transformed FTSEC can colonize the ovary, we developed an ovary-tumor co-culture system (Figure 7A). Our goal was to mimic the *in vivo* situation where the close proximity of the FT fimbria to the ovarian surface would increase the likelihood of early tubal HGSC precursors shedding, attaching, and invading the ovary. The ovary appears to provides an ideal microenvironment and scaffold for cancer cells to grow and expand before metastasizing to other organs (Perets et al. 2013; Yang-Hartwich et al. 2014). We labeled freshly isolated mouse ovaries with a green fluorescent dye and co-cultured the ovaries under ULA conditions with OVCAR8 or FTSEC that expressed either RFP or were labeled with a red fluorescent dye. After 24 hours of co-culture, we observed that for both OVCAR8 and OVCAR8-ΔL1 co-cultures, a small number of single cells attached to the ovary to a similar extent. However, between 2-7 days of co-culture, the majority of OVCAR8 cells expressing L1CAM that were not attached to the ovary, started to aggregate before attaching as spheres to the ovary surface (Figure 7A). These spheres then started to invade into the ovary (Figure 7B). In contrast, in OVCAR8-ΔL1 co-culture, the spheres attached to the ovary to a lesser extent and did not invade into the organ (Figure 7B, 7C).

**Figure 7:**
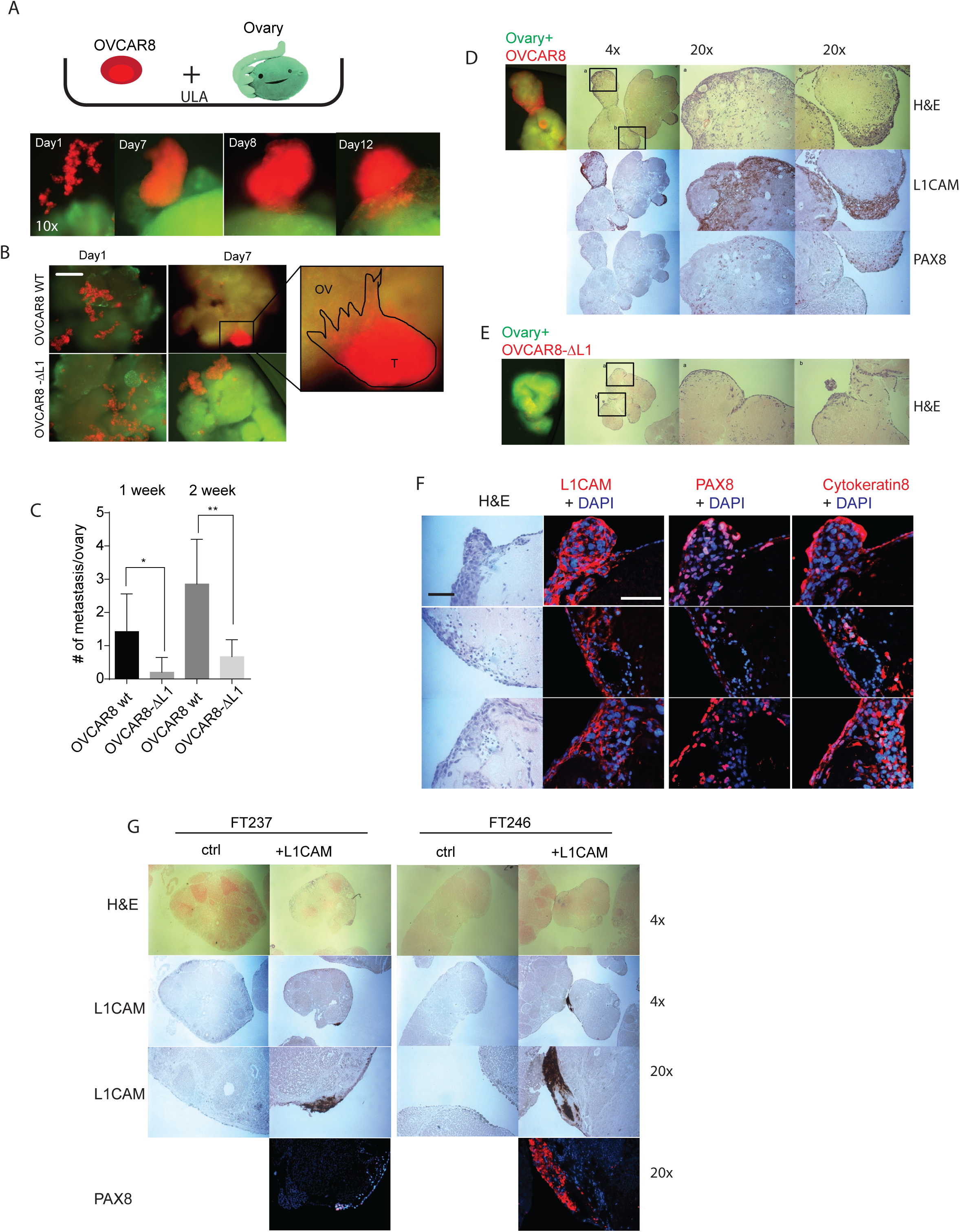
L1CAM is important for ovary invasion: (A) Upper row: Experiment schematic of co-culture of cells with an isolated mouse ovary under ULA culturing conditions. Lower row: representative observed invasion sequence over time. (B) Representative fluorescent image after one week’s co-culture of OVCAR8wt or OVCAR8-ΔL1 (red) with isolated ovary under ULA culture conditions. Scalebar represents 500 μm. (C) Bar graph representation of various OVCAR8 cells’ metastases to the ovary (more than 10 cells) after 1 and 2 weeks of co-culture. (D,E) Fluorescent and IHC images representing co-culture of OVCAR8wt or OVCAR8-ΔL1 with ovary, respectively. (F) fluorescent images of ovaries co-cultured with OVCAR8wt for expression of L1CAM, PAX8 and Cytokeratin 8. Scalebar represents 100 μm. (G) H&E and IHC images representing ovaries co-cultured with FT237 or FT246 transfected with L1CAM or a control vector.

We then analyzed co-cultures by IHC. We confirmed that the spheres of OVCAR8 attached and invaded into the ovary (Figure 7D). Interestingly, we also found invasion by OVCAR8-ΔL1 cells but it was mostly restricted to single cells and only at sites on the ovarian surface that exhibited a disruption of the ovarian surface epithelium that were formed by manipulation of the ovaries (Figure 7E). Furthermore, immunofluorescence of L1CAM, PAX8 and Cytokeratin 8 revealed that attached spheres of OVCAR8 cells spread both vertically below the OSE and laterally into the ovary tissue (Figure 7F).

Finally, we overexpressed L1CAM in FT237 and FT246 cell lines to examine whether L1CAM might also influence the ability of these cells to adhere and invade the ovary. The coculture of the FT cells with ovaries revealed an increased attachment and invasion of L1CAM-positive FT cells to the ovaries when compared to control cells, demonstrating the importance of L1CAM in metastatic attachment and invasion (Figure 7G). Overall, our results support the model whereby fallopian tube precursor cells utilize L1CAM to regulate integrin signaling to support ovarian cancer dissemination to the ovary.

## Discussion

Recent molecular studies of HSGC development (Eckert et al. 2016; Ducie et al. 2017; Labidi-Galy et al. 2017) emphasize the need to better understand the evolution of precursor FT lesions and factors that trigger metastatic dissemination to the ovary and beyond. Prior studies have shown that L1CAM, an essential cell adhesion molecule, is aberrantly expressed in tumor cells where it promotes cell migration and invasion (Schafer and Altevogt 2010; Kiefel et al. 2012). High levels of L1CAM are associated with poor overall survival in HGSC (Fogel et al. 2003; Bondong et al. 2012; Doberstein et al. 2014). Using immunohistochemistry for L1CAM on human FT precursor lesions, we found that peak L1CAM expression was largely restricted to cells that appeared to be detaching from the precursor lesion. Therefore, we hypothesized that L1CAM might have a functional role during the dissemination and metastasis of cells from STIC lesions to the ovary.

During the transition from STIC lesions to HGSC, metastasis to the ovary is thought to be a critical step (Perets et al. 2013; Yang-Hartwich et al. 2014). Presumably, the ovary provides a rich microenvironment and scaffold for rapid growth, expansion, and eventual dissemination. Yang-Hartwich et al. (2014) showed that the high content of collagen IV and the chemokine SDF-1 (CXCL12) are key factors to attract and maintain cancer cells in the ovaries. Additionally, using a mouse model, our lab showed that STIC lesions require spread to the ovary prior to peritoneal dissemintation (Perets et al. 2013). Importantly, the removal of ovaries in this mouse model resulted in a reduction of peritoneal metastasis, indicating the importance of the ovary as an optimal niche for cancer cells to grow and progress prior to becoming more widely metastatic. Cancer cells from STIC lesions have to master serial steps to successfully establish metastatic lesions in the ovary: 1) dissemination from the STIC lesion; 2) survival in the space between fallopian tube and ovary; 3) attachment to the ovary: 4) invasion into the ovary; and 5) establishing a self-sustained metastatic lesion.

We used a combination of approaches and models to characterize the role of L1CAM during these steps. We found that the ability of OVCAR8 to transition from a 2D monolayer to 3D spheres was abrogated by the loss of L1CAM. The loss of L1CAM also reduced the ability of cells to form spheres under ULA conditions which instead induced cell death. Conversely, overexpressing L1CAM in multiple normal FT cell lines led to an increase in sphere formation and reduced cell death under ULA conditions. These findings suggest that L1CAM enables survival of FT-derived cells under anchorage-independent conditions, thereby facilitating the dissemination step. The finding that L1CAM-specific antibodies were able to partly block the functions of L1CAM *in vitro* is consistent with animal studies showing that L1CAM-specific antibodies can block tumor progression in a peritoneal cell line xenograft model (Wolterink et al. 2010; Doberstein et al. 2015). Further studies are needed to establish whether L1CAM can be a target in this setting.

By co-culturing L1CAM-positive and -negative fallopian tube cells under ULA conditions, we showed that L1CAM-positive cells preferentially attached to other L1CAM-expressing cells. This might simulate the situation in a STIC lesion, in which the strong interaction of L1CAM-positive cells leads to the coalescence and dissemination of those cells. Indeed a number of studies have indicated that homotypic cluster formation can render cells resistant to anoikis (Hou et al. 2011; Iwanicki et al. 2016). Similarly, when co-culturing the OVCAR8 (expressing L1CAM) with FT cell lines under ULA conditions, the OVCAR8 cells assembled isolated colonies around the FT cells. In contrast, the OVCAR8-ΔL1 cells showed no preference to bind to each other and intermixed with the FT cells. Importantly, we found that L1CAM expression leads to the upregulation of Integrin-α5β1 and the recruitment of fibronectin to the basement membrane under 3D conditions, which subsequently led to the activation of the ERK and AKT pathways. The upregulation of Integrin-α5β1 has been shown to play an important role during tumorigenesis and in the promotion of metastasis in HGSC (Sawada et al. 2008; Deng et al. 2012; Kiefel et al. 2012; Iwanicki et al. 2016). Integrin-α5β1 has also been implicated in sphere formation and activation of the AKT and ERK pathways (Casey and Skubitz 2000; Casey et al. 2001; Ning et al. 2017). The recruitment of Integrin-α5β1 and fibronectin is important for the survival of cell aggregates that lose contact to their primary substratum, by helping to form anchorage to protect from anoikis (Yu et al. 2012; Serres et al. 2014).

For cancer cells to invade the ovary, cells have to detach from the FT, survive in the space between the FT and the ovary and attach to the OSE. To model and assess this process, we created an *ex vivo* system in which we cultured isolated ovaries together with cancer cells or fallopian tube cells under ULA culture conditions. Consistent with the findings from Yang-Hartwich et al, (2014) we observed that after 24 hours, some single cells attached to the ovarian surface at sites that showed a disruption of the OSE (Serres et al. 2014; Yang-Hartwich et al. 2014). During this first attachment stage, we did not observe differences between the L1CAM-positive or -negative cancer or FT cells. In general, we found that cells need to first coalesce into spheres before they can attach and invade the ovary. Interestingly, we observed that the spheres of L1CAM-positive cells were able to attach and invade through the OSE into the ovary stroma, whereas L1CAM-negative spheres were only able to attach weakly to the OSE and did not invade. Our data suggest that L1CAM enables survival of detached cells and promotes sphere formation that is a prerequisite for ovarian invasion. These L1CAM-dependent activities increase the likelihood to successfully establish a metastatic lesion in the ovary and lead to the ultimate progression to HGSC.

In summary, we examined the role of L1CAM in the early dissemination and survival of HGSC precursors from the FT. We showed that L1CAM is upregulated in precursor tumor cells and that it is required for sphere formation and survival under anchorage-independent conditions. L1CAM mediated these activities through upregulation of Integrin-α5β1 and fibronectin and activation of the AKT and ERK pathways. Our 3D *ex-vivo* model showed that L1CAM expression was important for FT cells to invade the ovary as a cohesive group. The genetic changes leading to L1CAM upregulation in this setting remain to be determined. The important role of L1CAM in the development and dissemination of HGSC suggests that L1CAM may have potential as a target for ovarian cancer prevention.

## Material and Methods

### Cell lines

The establishment of the fallopian tube cell lines has been previously described (Karst et al. 2011a; Karst and Drapkin 2012; Karst et al. 2014). They were cultured in Dulbecco’s Modified Eagle’s Medium (DMEM)/Ham’s F-12 1:1 (Cellgro) supplemented with 2% Ultroser G serum substitute (Pall Life Sciences, Ann Arbor, MI, USA) and 1% penicillin/streptomycin. All cancer cell lines (see Table 1) were cultured in RPMI 1640 (Invitrogen, Carlsbad, CA) supplemented with 10% fetal bovine serum (FBS, Atlanta Biologicals) and 1% penicillin/streptomycin (Invitrogen). All cells were grown at 37°C and a 5% CO_2_-containing atmosphere.

### Immunoblotting

Cells were lysed in RIPA buffer (25mM Tris-HCl pH 7–8, 150 mM NaCl, 0.1% SDS, 0.5% sodium deoxycholate, 1% Triton X-100 and protease inhibitors) for 3 h on ice. Protein content was quantified by BCA assay using the Pierce BCA kit protocol (#23228). 30 μgs of cell lysate were separated on a 4-15% gradient SDS-PAGE before being transferred to a PVDF membrane using the TurboBlot system (Bio-Rad). The membrane was blocked with 5% nonfat milk in TBST (Tris-buffered saline, 0.1% Tween 20) for one hour at room temperature. Blots were incubated in primary antibody (diluted in blocking buffer) overnight at 4°C. After washing, membranes were incubated in either anti-mouse or anti-rabbit HRP-conjugated secondary antibody (Cell Signaling, 1:1000) diluted in TBST for 1 hrs. Proteins were detected using Clarity Chemiluminescent HRP Antibody Detection Reagent (Bio-Rad, #1705061) and visualized with a ChemiDoc imaging system (Bio-Rad).

### TCGA analysis

Data from the TCGA database were extracted and downloaded from the XENA portal of the University of California, Santa Cruz (http://xena.ucsc.edu/). The extracted copy number and RNAseq data from the TCGA ovarian cancer cohort, TCGA pan-cancer cohort and the Cancer Cell Line Encyclopedia were analyzed with the GraphPad Prism or the ggplot2 package of the R software.

### Viability assay

Cell viability was monitored after 72 hrs, using a CellTiter 96® Non-Radioactive Cell Proliferation Assay (Promega, Mannheim, Germany). Each assay was performed in triplicate and repeated at least 3 times. Data are presented by means ± SD. Statistical and significant differences were determined by ANOVA with post-hoc analysis. Cells were additionally stained with crystal violet to count the remaining attached cells.

### Short-interfering RNA transfection

A pool of three short-interfering RNA (siRNA) duplexes of the trilencer-27 siRNA (Origene) was used to downregulate corresponding protein expression. As a negative control, a non-specific scrambled trilencer-27 siRNA was used. Twenty-four hrs. before transfection, 1 × 10^5^ cells were seeded in six-well plates (except OVCAR3, where 2 × 10^5^ cells were used). Transfection of siRNA was carried out using Lipofectamin RNAiMax (Invitrogen) together with 10 nM siRNA duplex per manufacturer’s instructions. To transfect 10 cm plates we scaled up our protocol by a factor of five. Conversely, to transfect one well of a 96-well plate we divided our protocol by 20.

### Immunofluorescence

For immunofluorescent analysis, cells were grown overnight in a 96-well Cell Imaging Plates (Eppendorf). Cells were then fixed in 4% (v/v) paraformaldehyde in PBS for 20 minutes at room temperature. Cells were blocked with super-block buffer (Thermo Scientific) and incubated with primary antibody overnight at 4°C. The secondary antibody was incubated for 1 hr. at room temperature. Detection was performed using secondary antibodies conjugated to Cy3 and Alexa 488 Fluor Dyes (Molecular Probes). Cells were then stained with DAPI and after an additional wash Flouromount-G (Sigma-Aldrich) was added. Cells were analyzed by microscopy using a Nikon E400 microscope.

### Immunohistology

Immunohistochemical staining was performed using Envision Plus/Horseradish Peroxidase system (DAKO, Carpinteria, CA, USA). Formalin-fixed paraffin-embedded tissue sections were de-waxed, rehydrated, and incubated in hydrogen peroxide solution for 30 min to block endogenous peroxidase activity. Antigen retrieval was carried out at 100°C treatment in citrate buffer (pH 6.0) for 20 min. Sections were incubated with primary antibody overnight at 4°C. The secondary antibody was applied for 30 min, followed by 3,3′-Diaminobenzidine (DAB) for 5 min.

### Migration assay

This assay was used to study the ability of cells to migrate through a coated transwell-plate placed (Corning) in a 24-well-plate. The bottom side of the polycarbonate membrane with a pore size of 5 μm was pre-coated with fibronectin (10 μg/ml) for 90 minutes at 37°C. The lower compartment was filled with 600 μL serum-free medium and the transwell insert was transferred into the well. The cells were diluted to 1×10^5^ cells in 100 μl in serum free medium and transferred to the upper compartment and incubated in a cell culture incubator for 16 h at 37°C. Cells in the upper compartment were removed using a cotton swab. Migrated cells were stained with 500 μl of a crystal violet buffer (Crystal Violet 0.05% w/v, Formaldehyde 1%, 10X PBS (1X), Methanol 1%) for 45 min. The transwell was washed extensively in water before the microporous membrane was cut out and transferred to a new well containing 300 μl 10% acetic acid. The eluted dye was transferred to a 96-well-plate and the absorption was measured in a plate reader at 570 nm.

### Ultra-low attachment assay

Cells were detached by 0.25% trypsin, resuspended in culturing media, centrifuged for 5 minutes at 1000 rpm and resuspended in serum free media. For the live/dead assay, 5×10^3^ cells were seeded per well of a 384-well ultra-low-adhesion (ULA) plate and 2 μM calcein, 3 μM of ethidium bromide was added to the cells and incubated for three hours. Cells were analyzed on a fluorescence microscope and mean intensity of the green and red fluorescence channel was analyzed with imageJ.

### Compaction assay

OVCAR8 cells were seeded in ULA 96-well round bottom plates (Corning). Cells were briefly spun at 127 g for 3 minutes. To generate cellular clusters, cells were returned to the incubator for 24 to 48 hours. Cellular clusters were then imaged for 48 hours at 20-to 30-minute intervals using a Nikon Ti-E inverted motorized widefield fluorescence microscope with integrated Perfect Focus System and (20×, 0.45 *numerical aperture* (*NA*)) magnification/NA phase/DIC optics equipped with a CO_2_-and temperature-controlled incubation chamber (Iwanicki et al. 2016).

### Colony formation assay

2000 cells were seeded on 6-well plates and cultured for two weeks. Cells were then washed with PBS and stained with the Crystal Violet buffer (Crystal Violet 0.05% w/v, Formaldehyde 1%, 10X PBS (1X), Methanol 1%) for 20 min. Cells were washed 5 times with water and air dried. Colonies were counted and size was calculated with ImageJ.

### CRISPR-Cas9 Targeting of L1CAM

To knock out the L1CAM gene in OVCAR8 cells, we used the GeneArt CRISPR system from ThermoFisher. Briefly, we co-transfected Cas9 mRNA with *in vitro* transcribed guide RNA (IVT gRNA) by using the transfection reagent MassengerMax. We targeted two sequences of L1CAM: Sequence 1: CAAAGCAGCGGTAGATGCCC.TGG

Sequence 2: TCATCACGGAACAGTCTCCA.CGG.

The transfected cells were then clonally separated and clones were screened for the loss of L1CAM expression.

### Overexpression

We transduced cells transiently with adenoviral particles (Vectorbuilder) generated from a 2^nd^ generation adenovirus gene expression vector containing either hL1CAM [ORF026248] under a CMV promoter or the control vector 72h before subsequent experiments were performed.

### 3D ovary invasion assay

Cancer or fallopian tube cells, cultured in a 10cm dish or flask, were detached with 0.25% trypsin, resuspended in culturing media, and centrifuged for 5 minutes at 1000 rpm. Cells not expressing a fluorescent marker were resuspended in serum free media and 1×10^5^ cells/ovary were incubated with a red fluorescent dye (CellTracker™ Red CMTPX, life technologies) for 30 minutes at 37°C before washing 3 times with PBS.

C57BL/6 mice were sacrificed by using a CO_2_ chamber before whole ovaries were harvested. All animal protocols were approved by the Institutional Animal Care and Use Committee (IACUC) at the Unversity of Pennsylvania. The ovaries were surgically separated from fallopian tubes and the bursa was removed. The ovaries were washed two times in PBS with 10% penicillin streptomycin on ice. Ovaries were then incubated with a green fluorescent dye (CellTracker™ Green CMFDA, life technologies) dissolved in serum free medium for 30 min at 37°C before washing 3 times with PBS. Organs and 1×10^5^ cells were then added to 1ml culturing media in an ultra-low attachment 24 well plate and imaged every 24 hrs. for up to 2 weeks for red and green fluorescence. All animal protocols were approved by the Institutional Animal Care and Use Committee at the University of Pennsylvania.

### 3D co-culture experiment

Cancer or fallopian tube cells, cultured in a 10cm dish or flask, were detached by 0.25% trypsin, resuspended in culturing media, and centrifuged for 5 minutes at 1000 rpm. Cells not expressing a fluorescent marker were resuspended in serum free media and incubated with a red or green fluorescent dye (CellTracker™ Red CMTPX, Green CMFDA, life technologies) for 30 minutes at 37°C before washing 3 times with PBS. Marked cell were co-cultured in normal media in a ratio of 1:10 of L1CAM expressing cells (red) to control cells (green), respectively.

### Statistics

All correlation values were calculated with the Pearson correlation coefficient. P-values were calculated with an unpaired two-sided t-test. *=p<0.05, **=p<0.01, ***=p<0.001, ****=p<0.0001

### Study approval

All human and animal protocols were approved by institutional review boards at the University of Pennsylvania.

## Acknowledgements

We thank members of the Drapkin lab for fruitful discussions and comments. We thank Paul Kroeger and Paul Whittaker for critical review of the manuscript and Teri Ord for assistance with the isolation of the mouse ovaries. This work was supported by the German Research Foundation DFG (K.D.), the Dr. Miriam and Sheldon G. Adelson Medical Research Foundation (R.D., G.B.M.), the Honorable Tina Brozman Foundation for Ovarian Cancer Research (R.D.), The Basser Center for BRCA (R.D.), the Claneil Foundation (R.D.), OCRFA (G.B.M), and NIH SPORE (1P50CA217685-01) in Ovarian Cancer (G.B.M).

## Author Contributions

Author contributions: K.D., and R.D designed research; K.D., R.S., S.S., M.P.I., M.F., L.E.S, K.M.D and Y.F. performed research; G.B.M. and P.A. contributed new reagents/analytic tools; K.D., R.D., and P.A. analyzed data; K.D. and R.D. wrote the paper, and all authors reviewed and edited the manuscript.

---

**Supplementary Figure 1:**
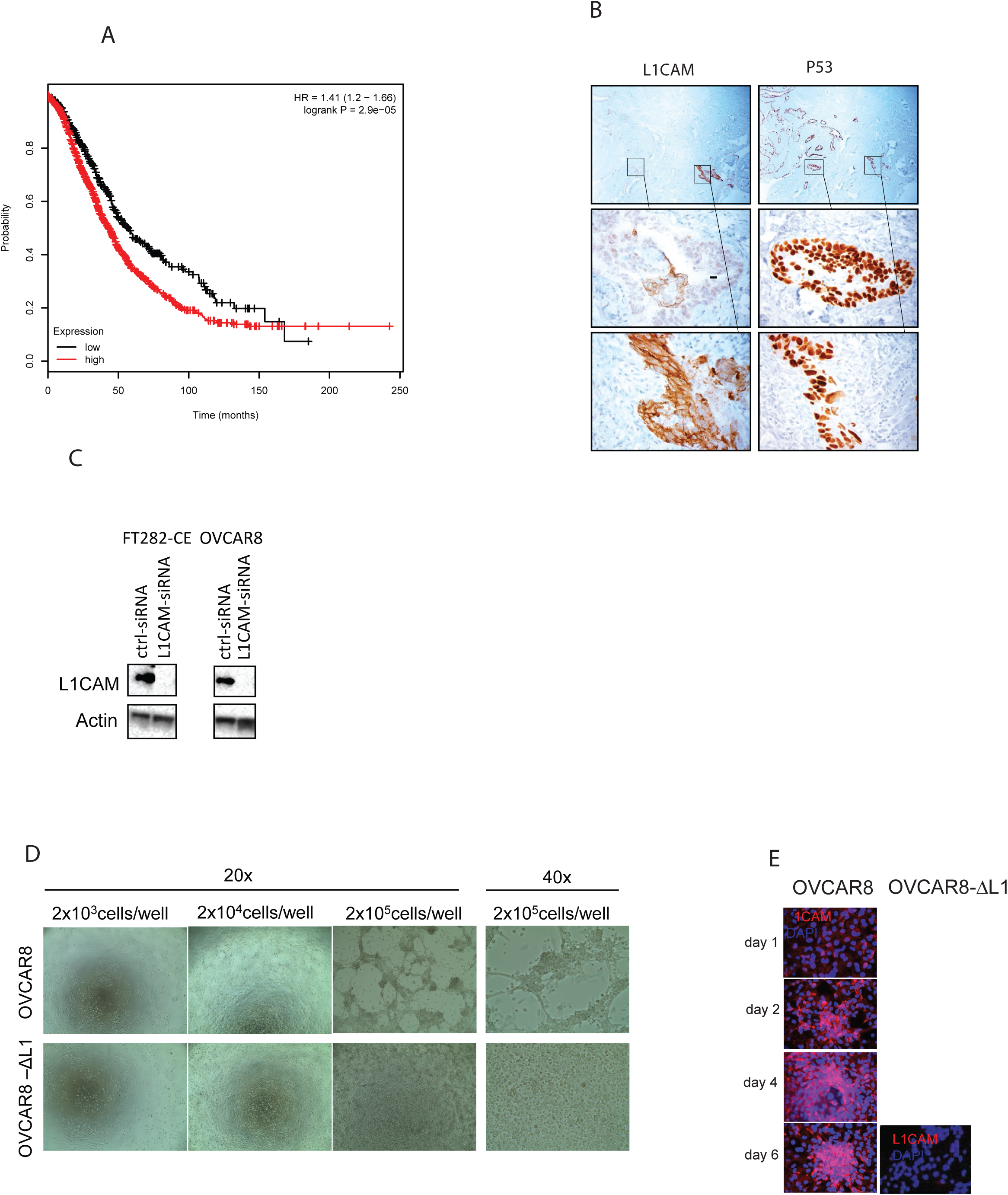
(A) Kaplan Meier overall survival analysis with the KM-Plotter of L1CAM (red line) against rest (black line). Cutoff value of 144 was used with the probe set 204584_at. n=1656, HR=1.44, p=6.1×10^−6^. (B) Upper panel: STIC lesion stained for L1CAM and P53. Magnified areas show areas of P53 positive but L1CAM low expression (middle panel) and areas that show a high L1CAM and P53 expression (lower panel). (C) Western blot analysis of FT282-CE and OVCAR8 after knock-down of L1CAM. (D) Bright filed picture of OVCAR8 (upper panel) andOVCAR8-ΔL1 (lower panel) seeded at increasing densities for 72 h in 24 well plates. (E) Fluorescent images of OVCAR8 and OVCAR8-ΔL1 representing L1CAM (red) and DNA (Dapi, blue) at different timepoints.

**Supplementary Figure 2:**
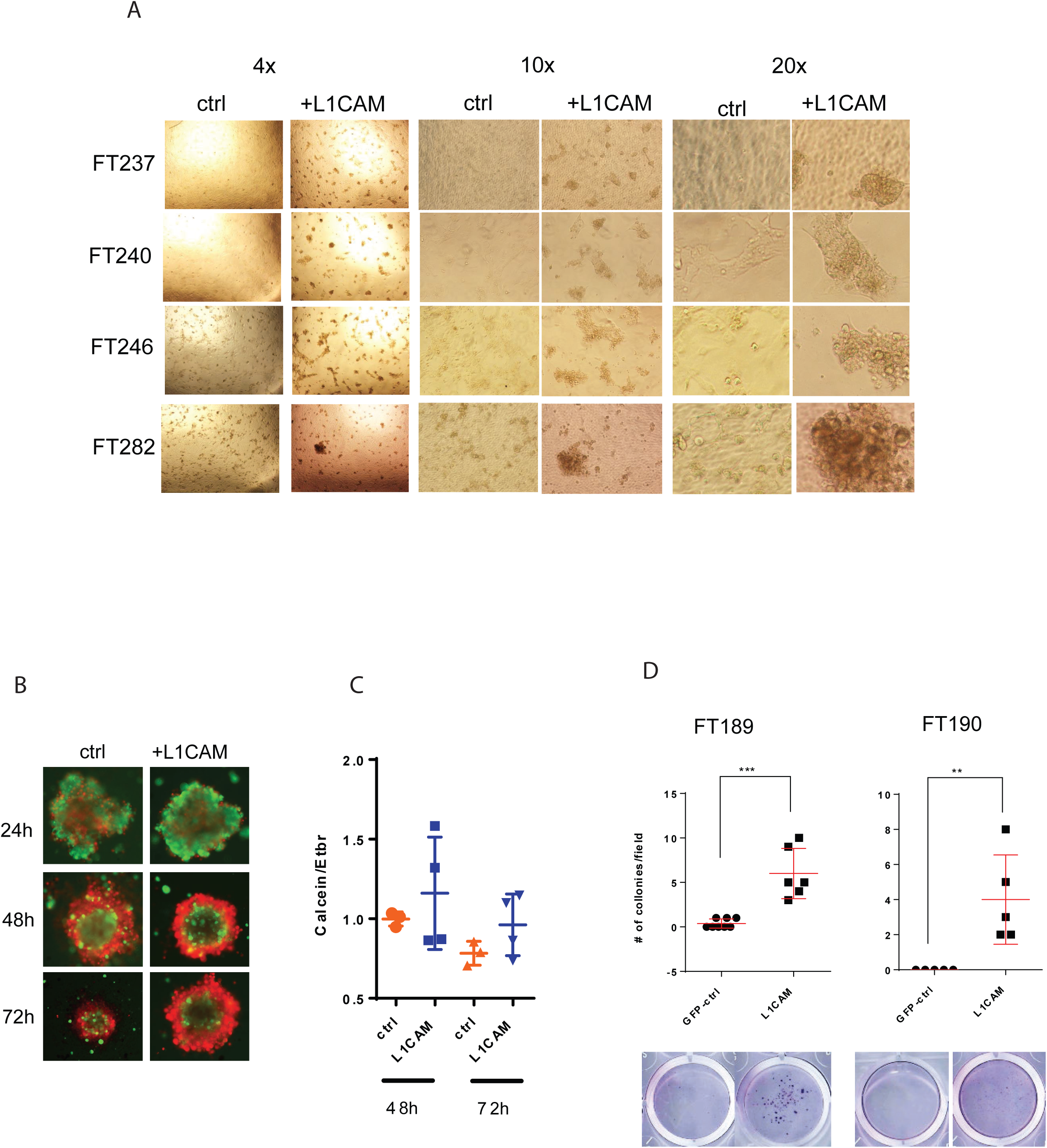
(A) Bright field analysis of FT237, FT240, FT246 and FT282 after transfection with either control or L1CAM vector. 24h after transfection, cells were detached and re-seeded on a new 24 well plate and grown for 48 h before pictures were taken. (B) Fluorescent images stained with calcein (green) and ethidium bromide (red) representing FT240 cells transfected with a L1CAM (right panel) or a control vector (left panel). (C) Dot plot depicting the ratio between living and dead cells, assessed by measuring the ratio between calcein (green) and ethidium bromide (red) intensity in FT240 cells transfected with L1CAM or a control vector after culturing under 3D and serum free conditions. (D) FT189 and FT190 were grown under ULA condition for 72 h before seeded to an adherent cell culture plate. Colonies were stained with crystal violet (lower panel) and colonies were counted (upper panel).

**Supplementary Figure 3:**
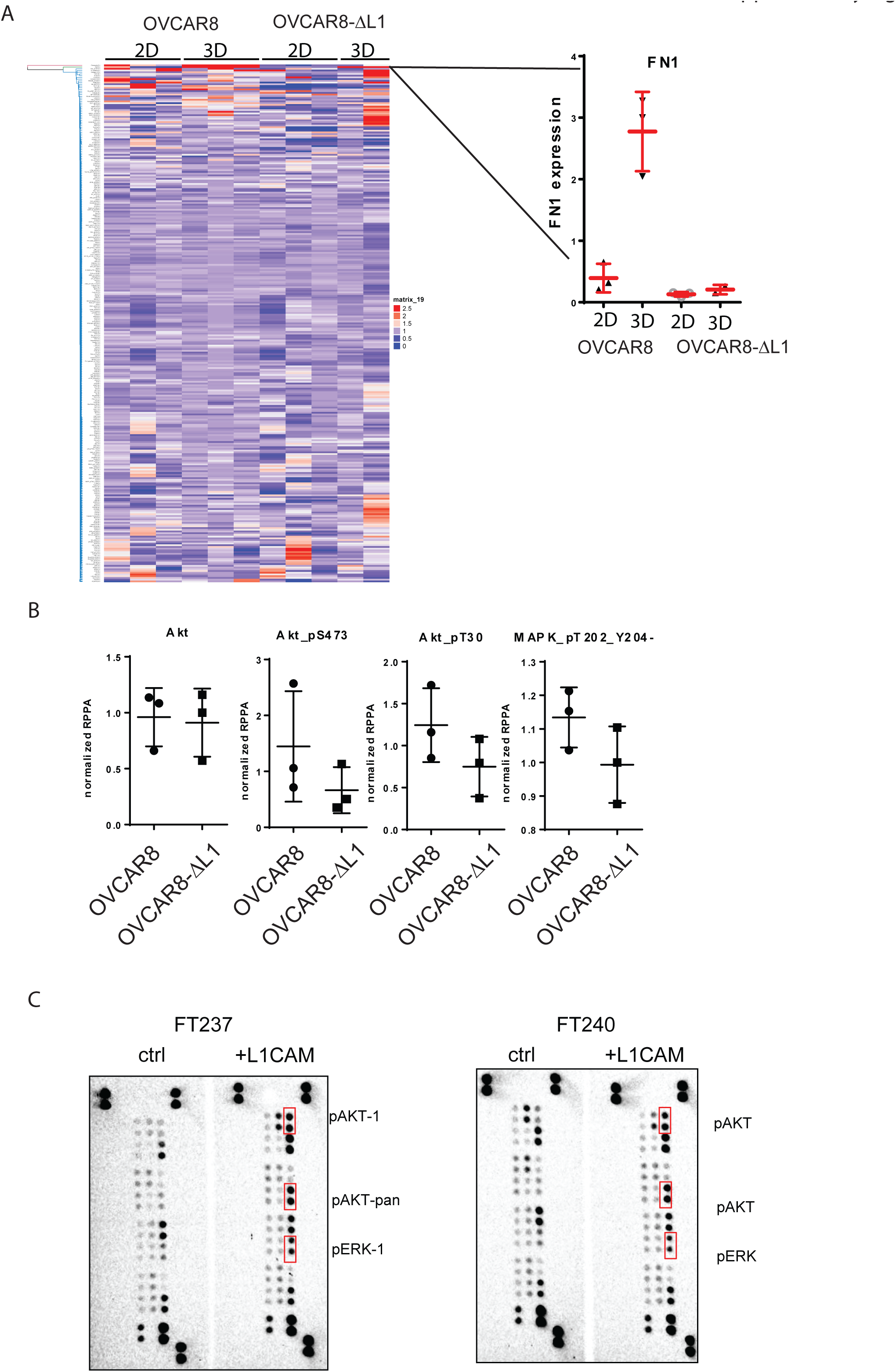
(A) RPPA analysis in OVCAR8 and OVCAR8-ΔL1 under 2D and 3D culturing conditions. (left) centroid clustered heatmap of samples. (right) Dot plot of fibronectin (FN1) expression from RPPA analysis. (B) RPPA analysis of AKT, AKT_pS473, AKT_pT30, and MAPK_pT02_Y204 in OVCAR8 and OVCAR8-ΔL1 cultured under 2D conditions. (C) MAPK arrays incubated with lysates of FT237 (left) or FT240 (right) transfected with a control or L1CAM vector.

**Supplementary Table 1.** List of cell lines used in these studies.

| Cell lines | cell line origins | Media |
| --- | --- | --- |
| Fallopian tube cell lines | Drapkin lab | DMEMF12+2% USG |
| DF-cell lines | Drapkin lab | DMEMF12+2% USG |
| TOV21G | ATCC | RPMI +10%FBS |
| OVCAR4 | NCI/DCTD | RPMI +10%FBS |
| OVCAR8 | NCI/DCTD | RPMI +10%FBS |
| COV318 | ATCC | RPMI +10%FBS |
| OVSAHO | Gottfried Konecny | RPMI +10%FBS |
| CAOV3 | ATCC | RPMI +10%FBS |
| Kuramochi | Gottfried Konecny | RPMI +10%FBS |
| CAOV4 | ATCC | RPMI +10%FBS |
| JHOS-2 | RIKEN | RPMI +10%FBS |
| JHOS-4 | RIKEN | RPMI +10%FBS |
| OVKATE | HSRRB | RPMI +10%FBS |

**Supplementary Table 2:**
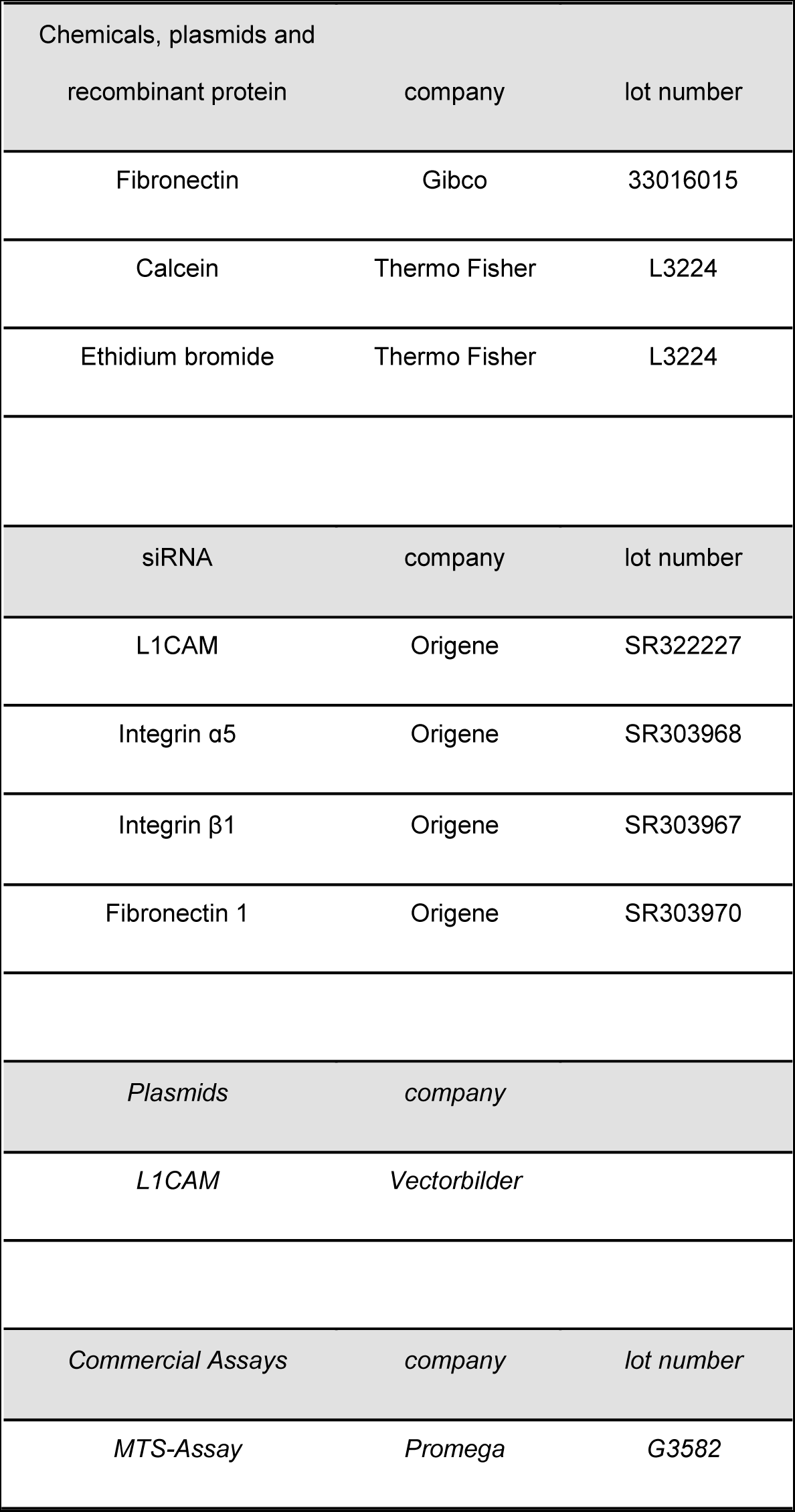
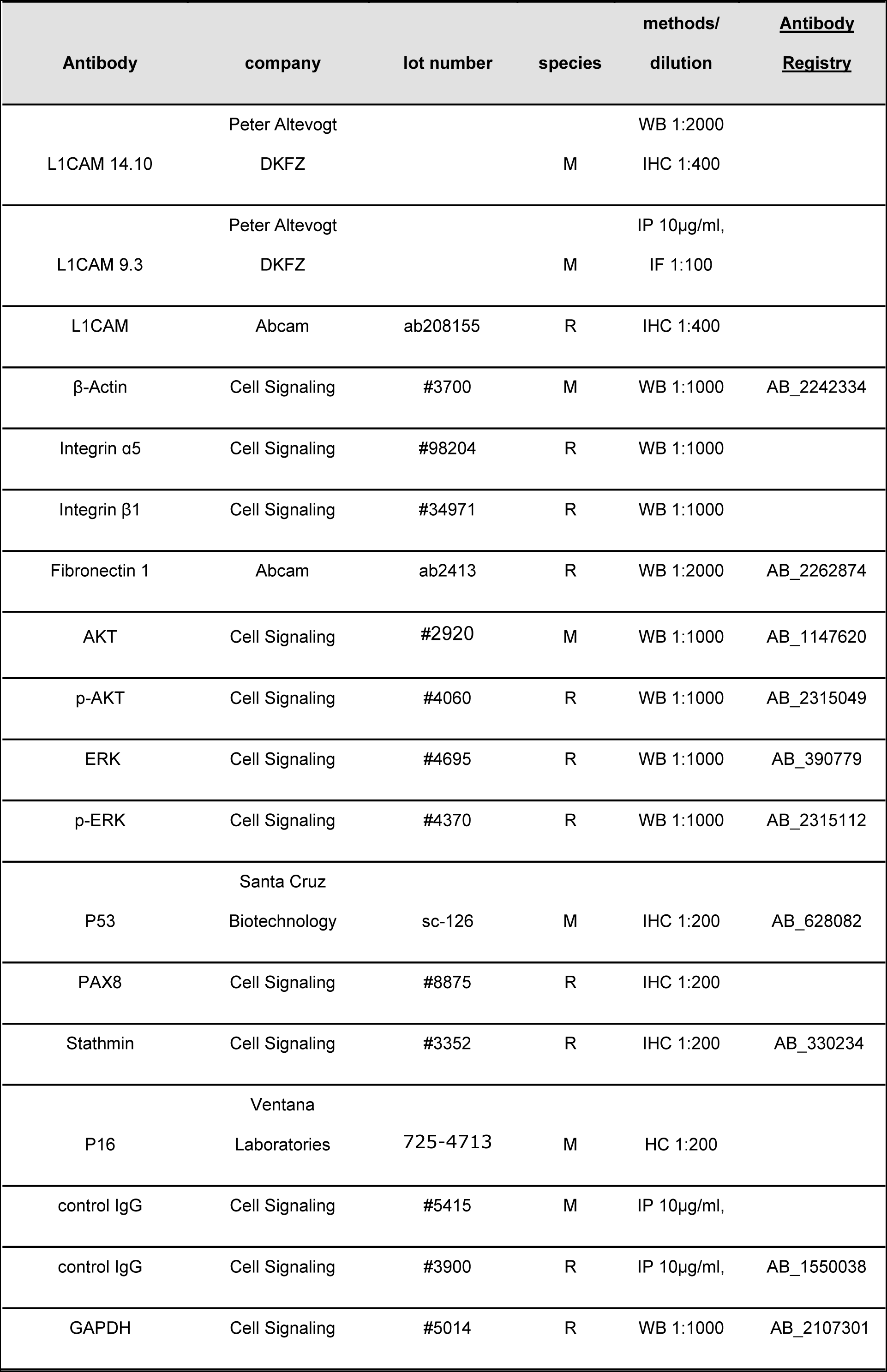
Materials

